# The CXCL12gamma chemokine immobilized by heparan sulfate on stromal niche cells controls adhesion and mediates drug resistance in multiple myeloma

**DOI:** 10.1101/2020.09.29.318303

**Authors:** Zemin Ren, Hildo Lantermans, Annemieke Kuil, Willem Kraan, Fernando Arenzana-Seisdedos, Marie José Kersten, Marcel Spaargaren, Steven T. Pals

**Author notes:** The authors share the last authorship (Marcel Spaargaren, Steven T. Pals). Correspondence to: Steven T. Pals, MD, Ph.D. Department of Pathology, Amsterdam University Medical Centers, Loc. AMC, Meibergdreef 9, 1105 AZ Amsterdam, The Netherlands.

## Abstract

The homing/retention, survival and proliferation of multiple myeloma (MM) cells critically depends on interaction with CXCL12 expressing stromal cells in the bone marrow (BM) niche. Here, we report a unique role in this interaction for the recently characterized CXCL12gamma isoform, which contains an extended C-terminal domain that binds heparan-sulfate proteoglycans (HSPGs) with an extraordinary high affinity. We observed that CXCL12γ is expressed *in situ* by reticular stromal cells in both normal and MM BM, as well as by primary BM stromal-cell (BMSC) isolates and BMSC lines. Importantly, upon secretion, CXCL12γ, unlike the CXCL12α isoform, was retained on the surface of these BMSCs. This membrane retention of CXCL12γ is HSPG-mediated, since it was completely annulated by CRISPR-Cas9 mediated deletion of the heparan-sulfate (HS) co-polymerase EXT1. Recombinant CXCL12γ was found to induce strong adhesion of MM cells to vascular cell-adhesion molecule 1 (VCAM-1) coated plates. Furthermore, CXCL12γ expressed by BMSCs and membrane-retained by HSPGs, supported robust adhesion of MM cells to the BMSCs. Specific genetic deletion of either CXCL12γ or of EXT1 significantly attenuated the ability of BMSCs to support MM cell adhesion and, in addition, impaired their capacity to protect MM cells from bortezomib-induced cell death. Our data indicate that CXCL12γ functions as a membrane-bound ‘niche chemokine’, which plays a unique role in the interaction of MM cells with the stromal niche by controlling adhesion/retention as well as cell adhesion-mediated drug resistance (CAM-DR). These findings designate CXCL12γ and associated HSPGs as potential therapeutic targets in MM.

## Introduction

The uncontrolled growth of cancer cells is driven by mutations in essential growth control genes, but their growth and survival is also strongly dependent on signals from the tumor microenvironment. In multiple myeloma (MM), a clonal expansion of malignant plasma cells in the bone marrow (BM), the interaction with specific BM-niches plays an important role in tumor-cell proliferation and survival. This interaction involves signaling via cell-surface receptors, including adhesion molecules, as well as by soluble factors secreted by various cells in the BM niche^1–3^. Despite improved survival due to the introduction of proteasome inhibitors, immunomodulatory drugs, and, more recently, monoclonal antibodies targeting MM cells^4–6^, MM is generally still incurable, which is largely due to the development of therapy resistance. MM cell interaction with the BM niche is believed to play a key role in this resistance, hence targeting this interaction presents a promising therapeutic strategy^1,7,8^.

The homing of hematopoietic stem cells (HSCs) as well as plasma-cell precursors to the BM is controlled by the chemokine CXCL12^9,10^. This chemokine also regulates the adhesion, transendothelial migration, and homing of MM cells to the BM by binding its receptor CXCR4 on the MM cells^11–13^. In the BM microenvironment, CXCL12 is mainly produced by specialized reticular BMSCs, also referred to as ‘CXCL12 abundant reticular (CAR)’ cells. Several splice variants of CXCL12 have been identified^14^, which all contain the CXCR4 binding motif but are differentially expressed in various murine and human tissues^15^. To date, the functional differences and biological significance of these distinct isoforms has remained largely unexplored. Virtually all *in vitro* functional studies, including those on MM cell migration and adhesion, have exclusively employed the CXCL12α isoform. Moreover, reported *in vivo* studies do not allow conclusions concerning the specific functions of the distinct CXCL12 isoforms, since the mice employed either carried a full deletion of CXCL12 or a deletion of CXCR4, the cognate receptor for all isoforms^16–19^. Interestingly, the recently characterized gamma-isoform of CXCL12 (CXCL12γ) has been shown to promote leukocyte accumulation and angiogenesis with a much higher efficacy than the ‘canonical’ CXCL12α isoform^15^. This enhanced biological activity of CXCL12γ is mediated by its extended C-terminal domain, which binds heparan sulfate proteoglycans (HSPGs) with an unprecedentedly high affinity^15,20^. Notably, in mouse BM, CXCL12γ was reported to be the dominant CXCL12 isoform. Furthermore, mice with a partial deletion in the HSPG-binding motives of CXCL12 showed increased numbers of circulating HSCs, suggesting a role for CXCL12-HSPG interaction in the retention of HSCs in the BM^21^.

HSPGs are membrane-bound or extracellular matrix proteins, consisting of a core protein decorated by covalently linked HS side-chains composed of repeating disaccharide units. These HS-chains undergo complex enzymatic modifications, which determine their binding capacity and specificity^22,23^ for a wide variety of morphogens, growth factor, and chemokines, thereby controlling the spatial distribution and activity of these ligand^24–26^. Given these properties, HSPGs appear well equipped to act as organizers of growth and survival niches. Indeed, studies in *Drosophila* have shown a crucial role for HSPGs in the germ cell-as well as hematopoietic-stem cell niches, controlling the activity of bone morphogenetic proteins (BMPs)^27,28^. In addition, HSPGs are known to bind a variety of proteins like Wnts, fibroblast growth factor (FGF), Midkine, and CXCL12, involved in the control of intestinal, neural, and hematopoietic niches^24,25,29^.

The extraordinary high affinity of CXCL12γ for HS, and its strong expression in mouse BM, prompted us to hypothesize that CXCL12γ could have a specific role in the organization of BM niches, including the plasma/MM cell niche. To explore this notion, we investigated the expression of this CXCL12 isoform in human BM and studied its functional role in the interaction of MM cells with BMSCs cells.

## Materials and Methods

### Cell culture

The human multiple myeloma cell lines (HMCLs) XG-1, MM1.S and L363 were cultured as described previously^29^. For XG-1, medium was supplemented with 500 pg/mL IL-6 (Prospec, Rehovot, Israel). BMSC lines HS5 and HS27a were cultured in DMEM (Invitrogen Life Technologies, Breda, The Netherlands) with 10% FBS (Invitrogen Life Technologies), 100 μg/ml streptomycin, 100 units/ml penicillin (Sigma Aldrich, St Louis, USA), Bone marrow endothelial cell lines HBMEC60 and 4LHBMEC were cultured in EGM-2MV medium (Lonza, Geleen, The Netherlands). Primary MM cells and BMSCs were derived from MM patients diagnosed at the Amsterdam University Medical Centers, location AMC, Amsterdam, the Netherlands. This study was conducted and approved by the AMC Medical Committee on Human Experimentation. Informed consent was obtained in accordance with the Declaration of Helsinki.

### Cloning, transfection and transduction

pLenti-CRISPR-sgEXT1 was constructed by inserting sgRNA-*EXT1* (GACCCAAGCCTGCGACCACG) into pL-CRISPR.EFS.GFP (Addgene plasmid # 57818) as previously described^29^. pLenti-CRISPR-sgCXCL12γ were constructed by inserting sgRNA-CXCL12γ#1(TTTAACACTGGCCCGTGTAC) and sgRNA-CXCL12γ#2 (AACTGTGGTCCATCTCGAGG) into pL-CRISPR.EFS.GFP. pBABE-CXCL12α and pBABE-CXCL12γ were constructed by inserting CXCL12α or CXCL12γ cDNA containing C-terminally C9-tagged (TETSQVAPA) sequences into pBABE-puro (Addgene plasmid # 1764). Lentiviral and retroviral particle production and transduction were performed as described before^29^.

### RT-PCR and genomic DNA PCR

Total RNA was isolated using TRI reagent (Invitrogen Life Technologies) according to the manufacturer’s instructions and converted to cDNA using oligo-dT. PCRs were conducted using SensiFast (Bioline, London, UK) on the CFX384 RT-PCR detection system (Bio-Rad). Isoform-specific primers sequences and housekeeping gene primers are shown in the Supplement Table 1. Genomic DNA was isolated using QIAamp DNA kit according to the manufacturer’s instructions. PCR primers used to detect CXCL12γ deletion are: forward primer: TCCCCAGTGGGAATCAGGTT; reverse primer: CTGGAGCTCCCAGGCTATTC.

### Adhesion assays

CXCL12α and CXCL12γ-induced adhesion to VCAM-1 was performed as described previously^31^. For adhesion to BMSCs and BM endothelial cells, MM cells were added to 96 well plates with confluent BMSCs or BM endothelial cells expressing a GFP marker. MM cells were spun down for 30 seconds at 400 RPM and subsequently incubated for 20 minutes to allow adhesion of MM cells to BMSCs or BM endothelial cells. Non-adherent cells were removed by washing with RPMI containing 1% BSA. Adherent cells were detached by trypsin and quantified by flow cytometry.

### Co-culture assays

For the co-culture assays, BMSCs were seeded in 96-well plates one day in advance to allow cell attachment. MM cells were added and incubated for 2 hours. Subsequently, drugs were added at the indicated concentrations. After 3 days, cells were collected and analyzed by flow cytometry, using 7-AAD (Thermo Fisher Scientific, Landsmeer, The Netherlands) to exclude dead cells. In the transwell assay, BMSCs were seeded in the lower compartment, and MM cells in the transwell insert (Costar, 0.4 μm; Corning, USA). After culturing the cells for 3 days in the presence or absence of bortezomib, the cell viability was analyzed by flow cytometry.

### Cell surface protein staining

Staining for HS was performed as described before^29^. Heparitinase used for digestion of cell-surface HS was purchased from amsbio (Abingdon, UK). For CXCL12γ cell-surface staining, cells were detached by 2μM EDTA, stained with isotype-specific antibody 6E9. Primary antibody binding was detected with rabbit anti-mouse IgG1-APC (Southern Biotech, Birmingham, USA). To assess binding of recombinant CXCL12γ, the HMCL XG1 was incubated with 1μg/ml recombinant CXCL12γ at 4°C for 90 minutes. After washing three times, cells were stained with mAb 6E9, binding was detected with rabbit anti-mouse IgG1-APC (Southern Biotech) and analyzed by flow cytometry.

### Immunohistochemistry

Paraffin-embedded BM biopsies for immunohistochemical studies were obtained from the Department of Pathology, Academic Medical Center, Amsterdam, the Netherlands. 4 μm tissue sections were treated with Tris-EDTA at pH 9 for 20 minutes at 121°C for antigen retrieval. Sections were incubated overnight at 4°C with the CXCL12γ isoform-specific mAb 6E9. Subsequently, the tissues were washed with PBS and incubated with rabbit-anti-mouse antibody (Southern Biotech) for 30 minutes at room temperature followed by poly-HRP-anti-rabbit IgG (DPVR110HRP, Immunologic, Duiven, the Netherlands) and Ultra DAB (Immunologic).

## Results

### CXCL12γ is expressed by human BM reticular stromal cells

CXCL12 produced by specialized, CAR-like, BMSCs cells has been shown to mediate the homing of both HSCs, plasmablasts, and MM cells to the BM^9,12^. However, to date, the expression of specific CXCL12 isoforms in the human BM microenvironment, and their possibly distinctive roles in the interaction with MM cells, has remained unexplored. To study CXCL12γ expression in human BM *in situ*, we employed immunohistochemistry, using mAb 6E9 specific for this isoform^15^. Interestingly, the CXCL12γ positive cells identified were reticular stromal cells with long cytoplasmic processes, which were scattered among hematopoietic cells, around adipocytes and capillaries (Figure 1A), areas with putative niche functions^32–37^. In BM samples infiltrated by MM cells, ample expression of CXCL12γ on stromal cells was also observed (Figure 1A).

**Figure 1.**
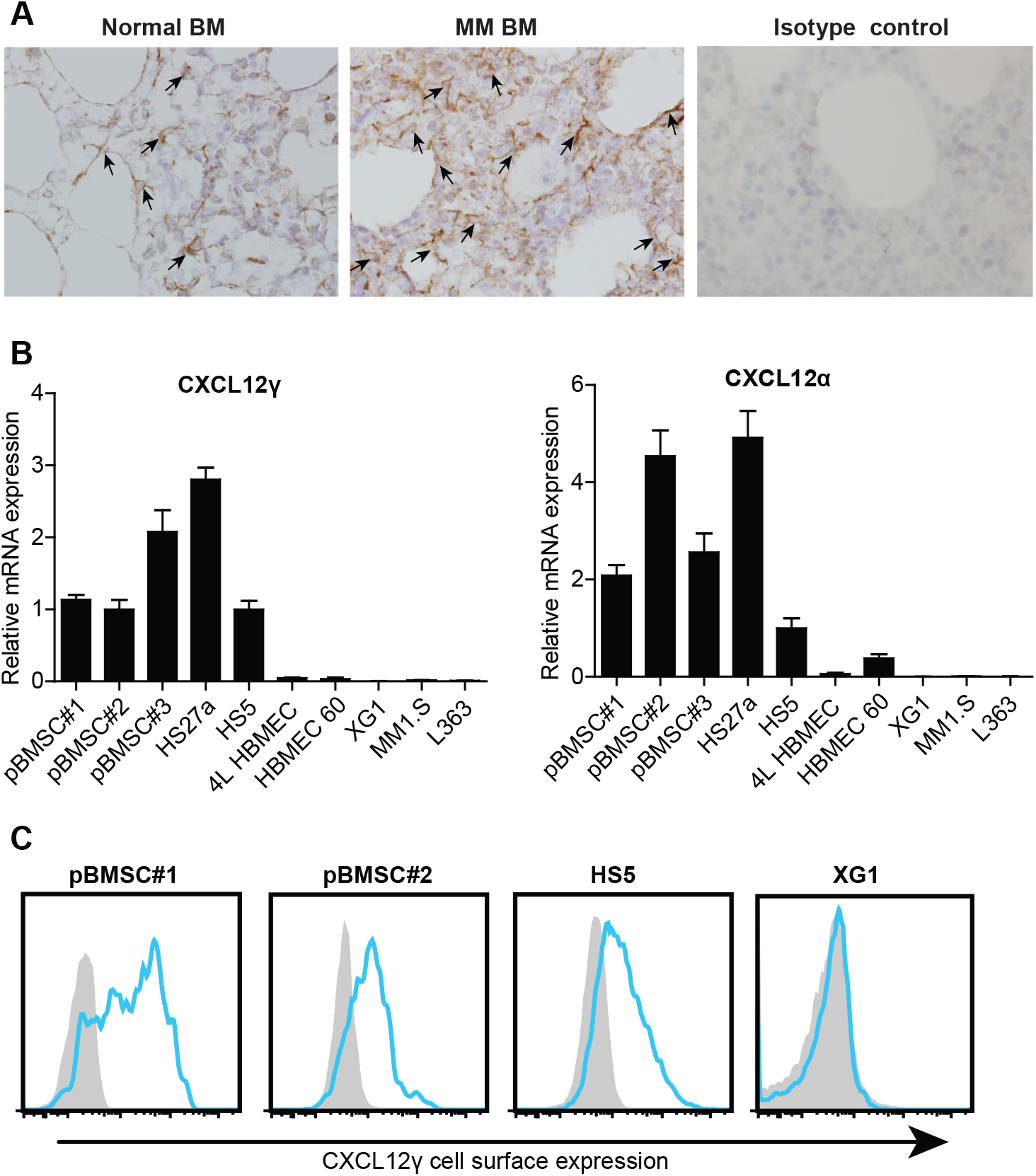
CXCL12γ expression by BMSCs. **(A)** CXCL12γ expression in normal BM, MM patient BM, as determined by immunohistochemical staining using the CXCL12γ isoform-specific mAb 6E9 (original magnification 200x). Arrowheads indicate CXCL12γ expression cells. **(B)** (left) CXCL12γ and (right) CXCL12α mRNA expression in the primary BMSC (pBMSC) samples, the human BMSC lines HS27a and HS5, the HBMEC lines 4LHBMEC and HBMEC 60, and the HMCLs XG1, MM1.S and L363, analyzed by qPCR and normalized to the housekeeping gene RPLPO. The mean ±SD of three independent experiments in triplicate is shown. **(C)** Cell-surface CXCL12γ expression on 2 primary BMSCs, BMSC line HS5, and the HMCL XG1, as determined by flow cytometry using mAb 6E9.

To define the expression of CXCL12γ by distinct BM-derived stromal cell types, we studied primary human BMSCs, the BMSC-lines HS5 and HS27, and the human BM endothelial cell (HBMEC) lines 4L-HBMEC and HBMEC60. Furthermore, we assessed CXCL12γ expression in various HMCLs, including XG1, MM1.S and L363. As shown in Figure 1B, primary BMSCs as well as BMSC lines were found to express both CXCL12γ and CXCL12α mRNA. By contrast, expression of both these CXCL12 isoforms was either low or undetectable in the HBMEC lines and in the HMCLs.

### CXCL12γ is immobilized on the cell surface of BMSCs by HSPGs

The C-terminal domain of CXCL12γ contains three positively charged HSPG-binding motives^15,20^. Exogenous overexpression of CXCL12γ in HEK293T cells has shown that this domain interacts with cell-surface expressed HSPGs, leading to immobilization on the cell membrane of the HEK293T cells^15^. We hypothesized that CXCL12γ expressed by BM reticular stromal cells might similarly be retained by HSPGs on the cell surface, and thereby function as a membrane-bound chemokine. Indeed, by employing the CXCL12γ-isoform-specific mAb 6E9, we observed that CXCL12γ is constitutively present on the cell-membrane of both primary BMSCs and the BMSC line HS5. No membrane-bound CXCL12γ was detected on the HMCL XG1, which does not express CXCL12γ mRNA (Figure 1C).

Both primary BMSCs and the BMSC line HS5 express high levels of cell-surface HSPGs, as detected by the HS-specific mAb 10E4 (Figure 2A). To study whether HS moieties indeed are responsible for the membrane-retention of CXCL12γ, we deleted *EXT1,* encoding the HS co-polymerase EXT1, which is critically required for the synthesis of HS-chains^29^. *EXT1* deletion in HS5 cells by CRISPR-Cas9 resulted in a complete loss of cell-surface HS expression, which was paralleled by loss of membrane-bound CXCL12γ (Figure 2B). Similarly, enzymatic removal of HS (Figure 2C-upper panel) from primary BMSCs by heparitinase resulted in a strong reduction of membrane-bound CXCL12γ (Figure 2C-lower panel). The HMCL XG1 expresses the HSPG syndecan-1^29^ but does not express endogenous CXCL12γ (Figure 1B and 1C). Incubation of XG1 cells with recombinant CXCL12γ resulted in strong membrane-binding, which was attenuated by *EXT1* deletion, corroborating the importance of HSPGs for CXCL12γ binding (Figure 2D).

**Figure 2.**
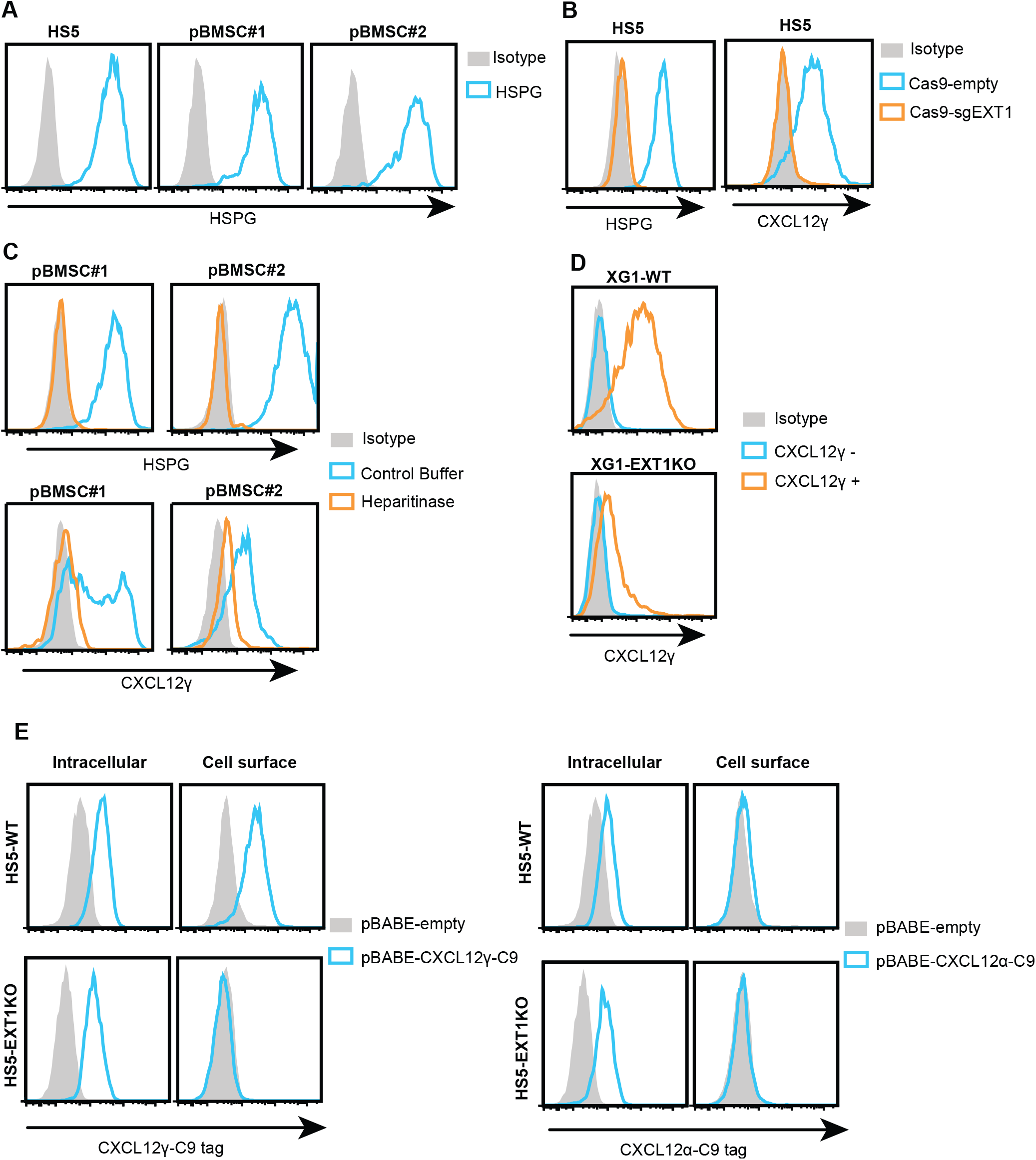
HSPGs retain CXCL12γ on the cell surface of BMSCs. **(A)** HSPG expression on the BMSC line HS5 and on primary BMSC samples, as determined by flow cytometry using mAb 10E4 against HSPG. **(B)** (left) HSPG cell-surface expression on HS5 cells transduced with either CRISPR sgEXT1 or CRISPR empty vector control; (right) CXCL12γ cell-surface expression on HS5 cells transduced with either CRISPR sgEXT1 or CRISPR empty vector control. **(C)** (upper panel) HSPG cell-surface expression on primary BMSCs treated with heparitinase or control buffer; (lower panel) CXCL12γ cell-surface expression on primary BMSCs treated with heparitinase or control buffer. **(D)** CXCL12γ cell-surface binding on XG1-WT (wild-type) or XG1-EXT1KO (knockout) cells after pre-incubation of the cells with 1 μg/ml recombinant CXCL12γ. **(E)** Exogenous cell-surface and intracellular CXCL12γ and CXCL12α expression. HS5-WT and HS5-EXT1KO cells were transduced with (left) pBABE-CXCL12γ-C9 or (right) pBABE-CXCL12α-C9. Intracellular and cell-surface expression of (left) CXCL12γ-C9 and (right) CXCL12α-C9 was detected with an anti-C9 antibody.

To assess whether membrane-retention indeed is a unique feature of CXCL12γ, not shared with the ‘canonical’ CXCL12α isoform, we expressed either C9-tagged CXCL12γ or CXCL12α in HS5 wild-type (WT) or HS5-EXT1KO cells. *Intracellular* expression of both isoforms was readily detected in both HS5-WT and HS5-EXT1KO cells (Figure 2E). However, *membrane-bound* CXCL12γ was only detected on the HS5-WT, but not on HS5-EXT1KO cells (Figure 2E-left), demonstrating that CXCL12γ requires HS for membrane retention. CXCL12α, by contrast, was not retained on the surface of either HS5-WT and HS5-EXT1KO cells (Figure 2E-right). These data demonstrate that CXCL12γ, unlike CXCL12α, is retained on the cell membrane of BMSCs by HSPGs.

### Recombinant CXCL12γ mediates MM cell adhesion to VCAM-1

It is well established that CXCL12α is able to induce VLA4-mediated adhesion of MM cells to VCAM-1^31,38^. To assess whether the γ-isoform of CXCL12 can similarly induce MM-cell adhesion, the HMCLs XG1, MM1.S and L363 were exposed to various concentration of either recombinant CXCL12α or CXCL12γ. As is shown in Figure 3A, both CXCL12α and CXCL12γ induced adhesion of these HMCLs to VCAM-1. The CXCL12α-induced adhesion showed a concentration-dependent bell-shaped curve, typical for chemokine/CXCL12α induced responses, with an optimum at 6.25 nmol. Remarkably, for CXCL12γ-induced adhesion this bell-shaped dose-response pattern was largely absent. At higher ligand concentrations, the CXCL12γ-induced adhesion was sustained and much stronger than the adhesion induced by CXCL12α. Notably, induction of MM cell adhesion required coating of CXCL12α and CXCL12γ to the adherence surface. In solution, both ligands were ineffective, indicating that CXCL12 immobilization is crucial for adhesion induction (Figure 3B).

**Figure 3.**
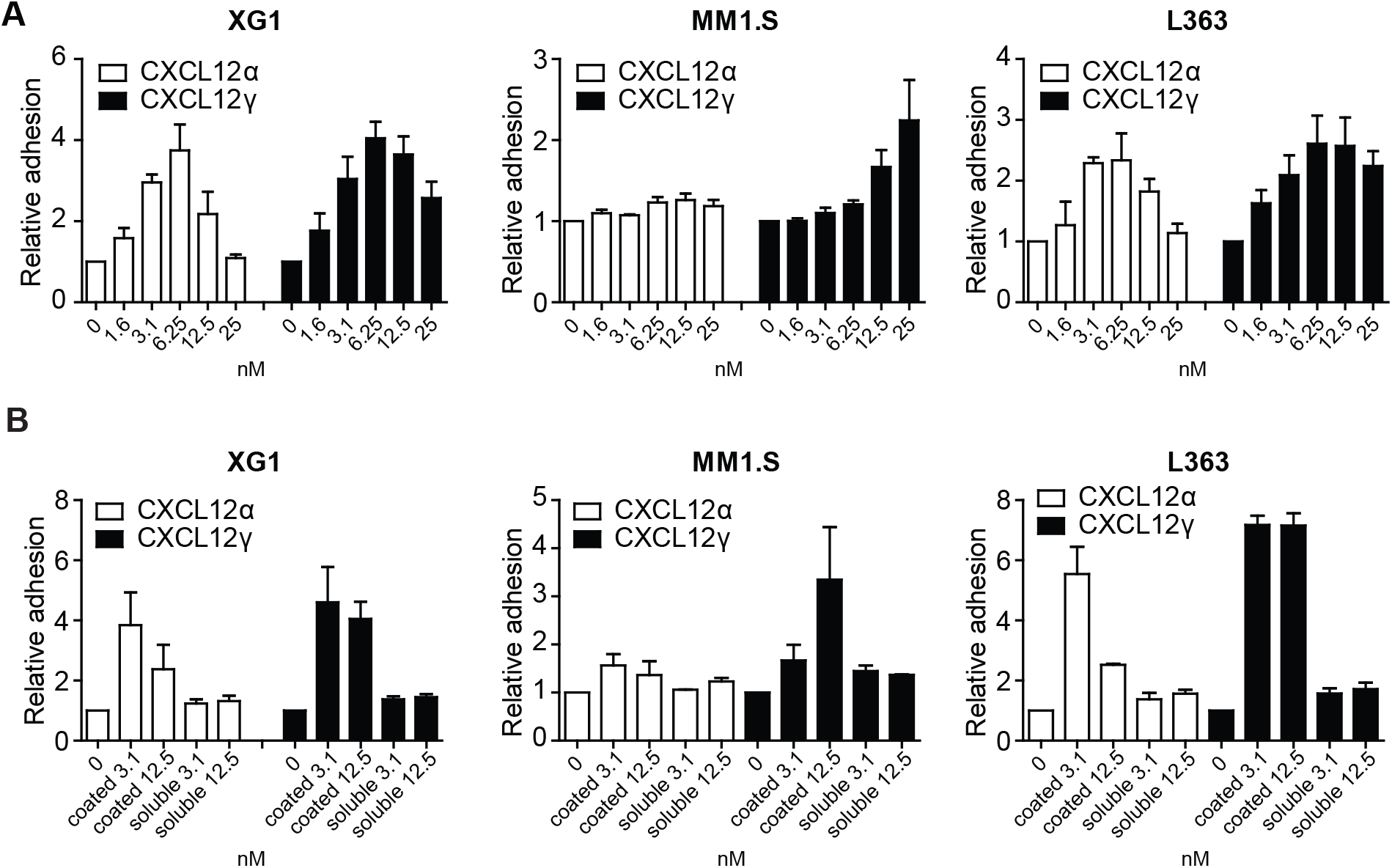
CXCL12α and CXCL12γ induce MM cell adhesion to VCAM-1. **(A)** Adhesion of XG1, MM1.S and L363 cells to a surface co-coated with either VCAM-1 and CXCL12α or CXCL12γ at indicated concentrations. Adhesion was allowed for 2 minutes and adhesion in the absence of CXCL12 was normalized to one. The mean ±SD of at least three independent experiments in triplicate is shown. **(B)** Adhesion of XG1, MM1.S, and L363 cells to VCAM-1. CXCL12α or CXCL12γ at indicated concentrations (nM) were either co-coated with VCAM-1 (coated) or present in the medium in soluble form (soluble). Adhesion in the absence of CXCL12 was normalized to one. Mean ±SD of at least three independent experiments in triplicate is shown.

### CXCL12γ expressed and membrane-retained by HSPGs on reticular stromal cells, mediates adhesion of MM cells

Given our finding that CXCL12γ is expressed and membrane retained by stromal niche cells, we hypothesized that this isoform might play a specific role in controlling MM adhesion to and retention in the BM niche. To specifically study the biological function of CXCL12γ, we employed two CRISPR sgRNAs designed to target the CXCL12 gene upstream and downstream of the fourth exon encoding the unique C-terminal tail of CXCL12γ (Figure 4A). Deletion in HS5 cells yielded a PCR product with a predicted size of approximately 500bp (Figure 4A) and was verified by Sanger sequencing (Figure 4B). Moreover, deletion was confirmed by loss of cell surface CXCL12γ protein expression (Figure 4C). Importantly, as anticipated, the expression of CXCL12α was not affected by deletion of exon 4 of the CXCL12 gene (Supplemental Figure 1). Furthermore, deletion of CXCL12γ had no effect on BMSC growth (Supplemental Figure 2).

**Figure 4.**
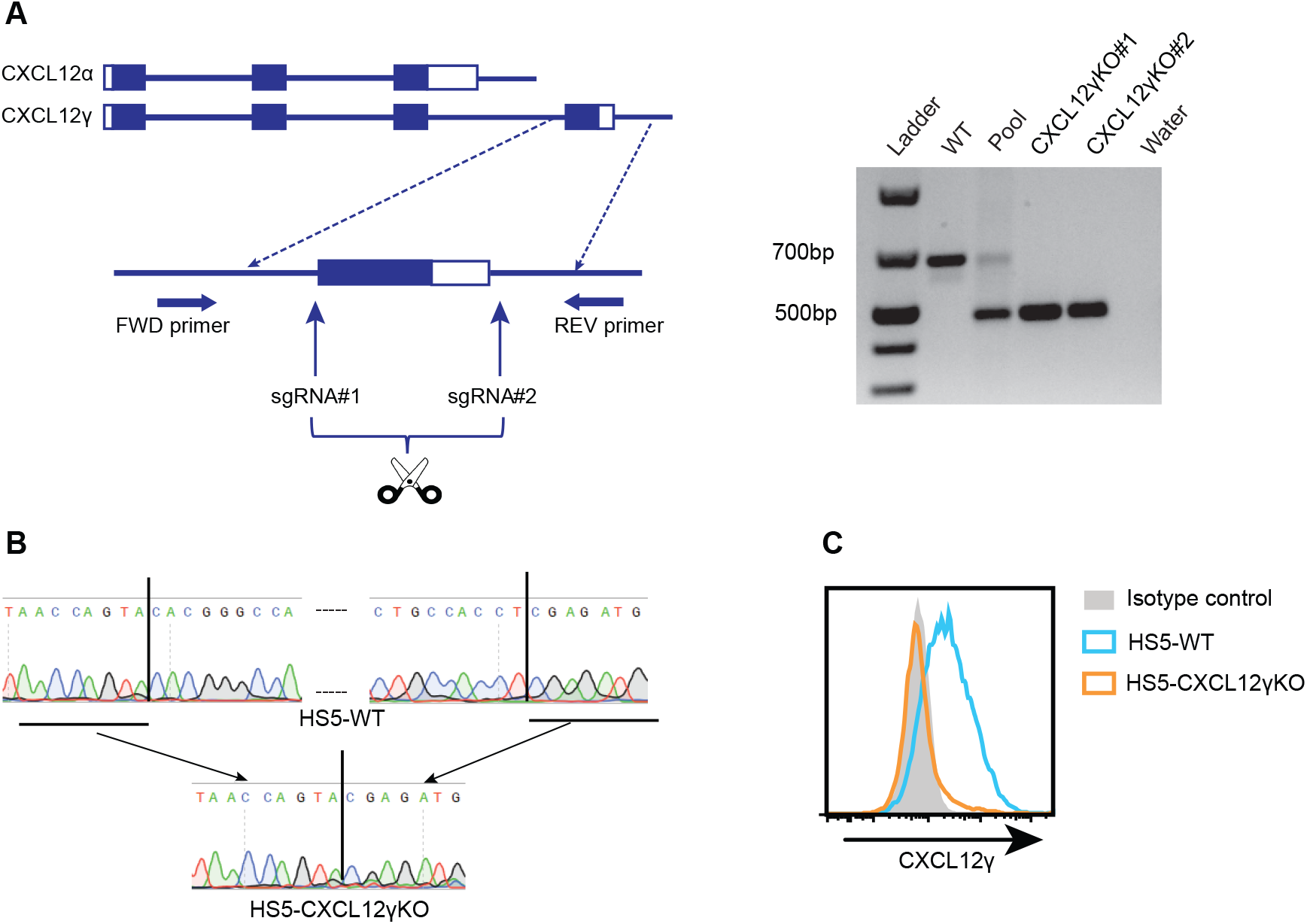
Isoform-specific dual-sgRNA CRISPR-mediated CXCL12γ KO. **(A)** Schematic representation of the CRISPR-induced deletion of the fourth exon (C-terminal tail) of CXCL12γ. sgRNA#1 was designed to target upstream of the fourth exon of CXCL12γ, and sgRNA#2 was designed to target the 3’UTR of the fourth exon. Rectangles represent exons and filled rectangles represent coding sequences. Lines indicate introns. (right) PCR analysis of the deletion of CXCL12γ; primers used are as indicated in panel A. Genomic DNA was isolated from HS5 cells with the CRISPR empty vector (WT); or co-transduced with CRISPR sgRNA#1 and CRISPR sgRNA#2, either before single-cell cloning (pool) or from two single cell KO clones (CXCL12γKO#1 and #2). Water was used as negative control. DNA ladder size is indicated on the left. **(B)** Confirmation of CRISPR-induced deletion by Sanger sequencing. Genomic DNA was isolated from HS5 cells transduced with empty vector CRISPR (HS5-WT) or CRISPR sgRNA#1 and CRISPR sgRNA#2 (HS5-CXCL12γKO). The CRISPR cutting sites are indicated by vertical line in HS5-WT; **(C)** Confirmation of CXCL12γ deletion by flow cytometry. Cell surface expression of CXCL12γ on HS5 cells, transduced with either empty vector CRISPR (HS5-WT) or HS5-CXCL12γKO were stained with the CXCL12γ specific mAB 6E9.

As is shown in figure 5A, the HMCLs XG1 and MM1.S displayed strong adhesion to HS5 and HS27 BMSCs, but did not adhere to BMECs. Interestingly, the capacity of HS5-CXCL12γKO cells to support adhesion of these MM cells was significantly reduced (Figure 5B and Supplemental Figure 3A). Likewise, the HS5-EXT1KO cells, which no longer express the HS-moieties required for membrane retention of CXCL12γ, also displayed a reduced capacity to support adhesion of these MM cells (Figure 5C and Supplemental Figure 3B). Similar to the adhesion of HMCLs, primary MM cells also showed a reduced adhesion to HS5 BMSCs lacking either CXCL12γ or EXT1 (Figure 5D). Importantly, exogenous reconstitution of CXCL12γ, completely restored the adhesion defect in the HS5-CXCL12γKO BMSCs, confirming the role of CXCL12γ. By contrast, exogenous overexpression of CXCL12γ in HS5-EXT1KO BMSCs could not rescue the defective adhesion of MM cells to these cells (Figure 5E), confirming the critical role of HSPG-mediated CXCL12γ cell-surface retention. Taken together, these data indicate that CXCL12γ expressed by BMSCs and immobilized by cell-surface HSPGs plays an important role in mediating MM cell adhesion to BMSCs.

**Figure 5.**
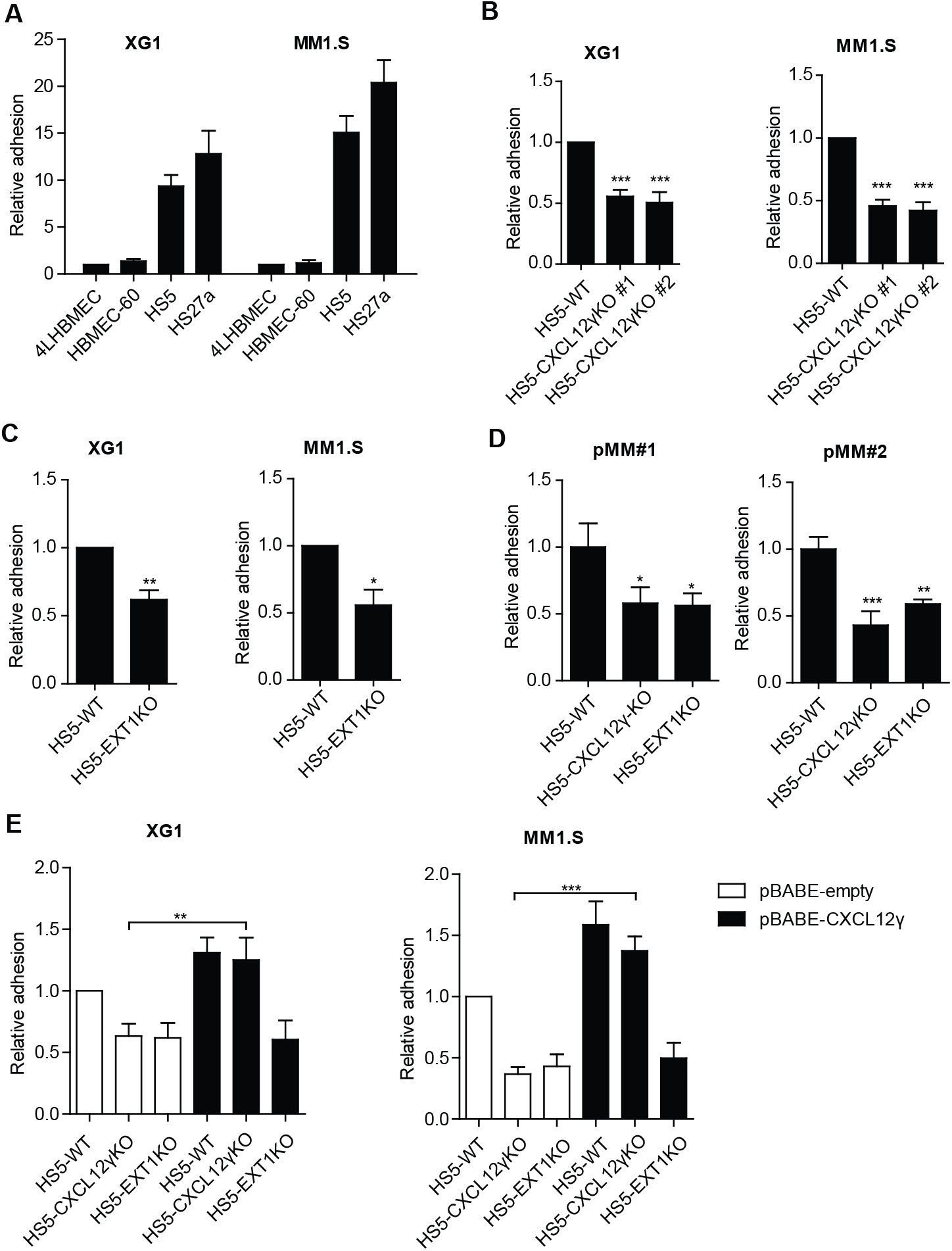
CXCL12γ retained by membrane-bound HSPGs promotes adhesion of MM cells to BMSCs. **(A)** Adhesion of the HMCLs XG1 and MM1.S to the BMSC lines HS5 and HS27a, and the HBMEC lines 4LHBMEC and HMMEC-60. Adhesion to 4LHBMEC is normalized to one. Mean ±SD of three independent experiments in triplicate is shown. **(B)** Adhesion of XG1 and MM1.S cells to HS5-WT and two independent CXCL12γKO clones. The mean ±SD of three independent experiments in triplicate is shown. Adhesion to HS5-WT is normalized to one. ***, P ≤ 0.001 using one-way ANOVA analysis. **(C)** Adhesion of XG1 and MM1.S cells to HS5-WT or HS5-EXT1KO cells. Mean ±SD of three independent experiments in triplicate is shown. Adhesion to HS5-WT is normalized to one. *, P ≤ 0.05; **, P ≤ 0.01 using unpaired student’s t-test. **(D)** Adhesion of primary MM cells from two patients to HS5-WT, HS5-CXCL12γKO, or HS5-EXT1KO cells. A representative plot for 2 independent experiments performed in triplicate is shown. *, P ≤ 0.05; **, P ≤ 0.01; ***, P ≤ 0.001 using one-way ANOVA analysis. **(E)** Adhesion of XG1 and MM1.S cells to HS5-WT, HS5-CXCL12γKO, or HS5-EXT1KO cells transduced with either pBABE-empty or pBABE-CXCL12γ vector. Mean ±SD of three independent experiments in triplicate is shown. **, P ≤ 0.01; ***, P ≤ 0.001 using one-way ANOVA analysis.

### CXCL12γ membrane-retained by HSPGs on BMSCs mediates resistance of MM cells to proteasome inhibitors

Interaction of MM cells with BMSCs plays a central role in the homing, retention, growth, and survival of MM cells as well as in drug resistance^1,2,40,41^. Inhibition of the CXCL12/CXCR4 axis, disrupting interaction of MM cells with BMSCs, has been reported to alleviate the protective effect of BMSCs, enhancing the sensitivity of MM cells to various drugs^1,8^. Given our observation that BMSC-derived CXCL12γ plays an important role in the adhesion of MM cells to BMSCs, we addressed the possible involvement of CXCL12γ in the protective effect of BMSCs against drug-induced MM cell death.

To measure MM cell death and the protective effect of BMSCs, the HMCLs XG1, MM1.S and L363 or primary MM (pMM) cells were co-cultured with HS5 BMSCs expressing green-fluorescent protein (GFP), to allow easy discrimination of both cell types (Supplemental Figure 4A). We focused on bortezomib since it represents a mainstay of current MM therapies. Moreover, unlike MM cells, which are highly sensitive, BMSCs are bortezomib insensitive *in vitro* (Supplemental Figure 4B), allowing reliable quantification of MM-specific cell death. As shown in Figure 6A-C, co-culture with HS5-WT BMSCs protected both HMCLs and pMMs from bortezomib-induced cell death. Interestingly, this protective effect was significantly reduced in MM cells co-cultured with HS5-CXCL12γKO cells, indicating involvement of CXCL12γ in mediating bortezomib resistance (Figure 6A and C).

**Figure 6.**
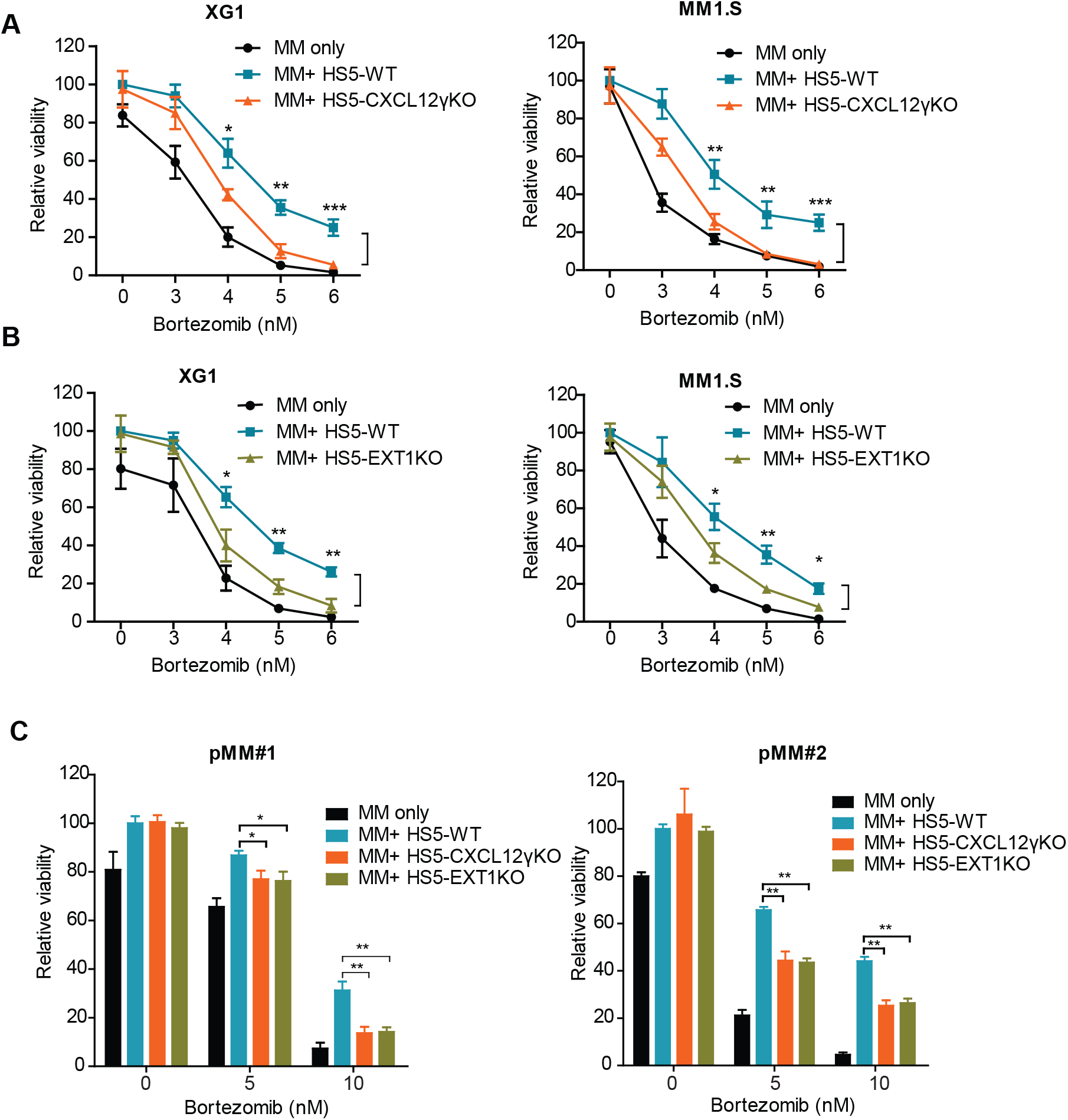
BMSCs provide bortezomib-resistance to MM cells through CXCL12γ and HSPGs. **(A)** HS5-CXCL12γKO cells show a reduced capacity to protect HMCLs from bortezomib-induced cell death. XG1 or MM1.S cells were cultured alone, or co-cultured with HS5-WT or HS5-CXCL12γKO BMSCs, in the presence of bortezomib at indicated concentrations for 3 days. The viability of the MM cells was analyzed by flow cytometry. Mean ±SD of three independent experiments in triplicate is shown. *, P ≤ 0.05; **, P ≤ 0.01; ***, P ≤ 0.001 using one-way ANOVA analysis; **(B)** HS5-EXT1KO cells show a reduced capacity to protect HMCLs from bortezomib-induced cell death. XG1 and MM1.S cells were cultured alone, or co-cultured with HS5-WT or HS5-EXT1KO BMSCs, in the presence of bortezomib at indicated concentrations for 3 days. The viability of the MM cells was analyzed by flow cytometry. Mean ±SD of three independent experiments in triplicate is shown. *, P ≤ 0.05; **, P ≤ 0.01 using one-way ANOVA analysis; **(C)** HS5-CXCL12γKO and HS5-EXT1KO cells show a reduced capacity to protect primary MM cells from bortezomib-induced cell death. Primary MM cells from two patients were cultured alone, or co-cultured with HS5-WT, HS5-CXCL12γKO or HS5-EXT1KO BMSCs, in the presence of bortezomib for 3 days. The viability of the MM cells was analyzed by flow cytometry. Plot representative for two independent experiments performed in triplicate. *, P ≤ 0.05; **, P ≤ 0.01 using one-way ANOVA analysis.

Since deletion of EXT1 results in loss of CXCL12γ membrane retention (Figure 2B), we examined if EXT1KO would also reduce the protective effect of BMSCs. Indeed, similar to HS5-CXCL12γKO cells, HS5-EXT1KO cells showed a significantly reduced capacity to protect both HMCLs and pMMs against bortezomib-induced cell death (Figure 6B and C). Similarly, the HS5-CXCL12γKO cells and HS5-EXT1KO cells also showed a reduced capacity to protect XG1 against cell death induced by carfilzomib, another commonly used proteasome inhibitor (Supplemental Figure 5). Recombinant CXCL12γ (or CXCL12α), in the absence of BMSCs, but co-coated with VCAM-1 did not affect the bortezomib sensitivity of HMCLs (Supplemental figure 6A and B). These findings indicate that the BMSC-mediated resistance to proteasome inhibitors involves CXCL12γ retained on the cell membrane of BMSCs by HSPGs.

### BMSC derived-CXCL12γ and HSPGs mediate CAM-DR

Drug resistance mediated by the MM BM microenvironment can either be caused by soluble factors or by direct physical cell-cell interactions mediated by cell adhesion molecules, termed soluble factor-mediated drug resistance (SFM-DR) and cell adhesion-mediated drug resistance (CAM-DR), respectively^1,42,43^. To directly investigate the cell-cell contact dependency of the BMSC-mediated resistance to bortezomib and establish if soluble factors released by BMSCs were (also) involved, we employed transwell co-cultures to physically separate MM cells from BMSCs. As shown in Figure 7A, in the transwell setting, HS5 BMSCs weakly, but significantly, protected the HMCL XG1, but not MM1.S, from bortezomib-induced cell death. This protective effect was not influenced by deletion of either CXCL12γ or EXT1. However, in a direct-contact setting, in which MM cells in suspension were removed before determining cell viability, BMSCs conferred a much stronger drug resistance to both XG1 and MM1.S. Importantly, this protective effect was largely abrogated by deletion of CXCL12γ or EXT1 and, hence, was CXCL12γ and HSPG-dependent (Figure 7B). Thus, CXCL12γ and HSPG on the cell surface of BMSCs promote MM cell adhesion to these BMSCs and thereby play an important role in CAM-DR.

**Figure 7.**
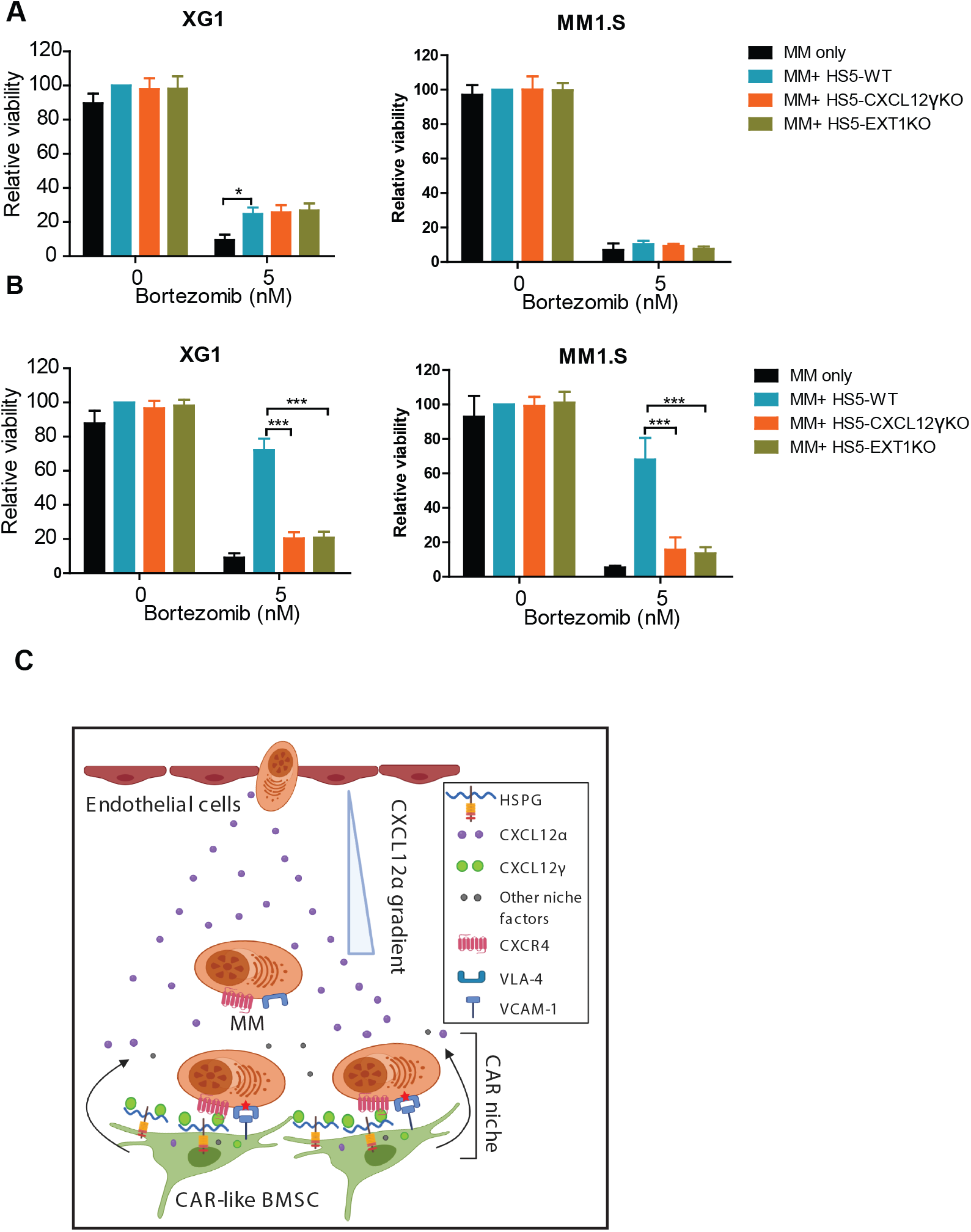
BMSC-derived CXCL12γ immobilized by HSPGs mediates CAM-DR. **(A)** Transwells co-culture of MM cells with BMSCs. XG1 and MM1.S cells were cultured either alone in the upper compartment or co-cultured with HS5-WT, HS5-CXCL12γKO or HS5-EXT1KO in the lower compartment of the transwells, in the presence of bortezomib for 3 days. So, the transwells physically separate the MM cells from the HS5 BMSCs. The viability of the MM cells was analyzed by flow cytometry. Mean ±SD of three independent experiments in triplicate. *, P ≤ 0.05 using one-way ANOVA analysis; **(B)** CXCL12γ-controlled drug resistance requires direct cell-cell contact between MM cells and BMSCs. MM cells were cultured, either alone or co-cultured with HS5-WT, HS5-CXCL12γKO or HS5-EXT1KO, in the presence of bortezomib for 3 days. The MM cells in suspension were removed. The viability of the remaining MM cells was analyzed by flow cytometry. Mean ±SD of three independent experiments in triplicate is shown. ***, P ≤ 0.001 using one-way ANOVA analysis. **(C)** Model for the role of distinct CXCL12 isoforms in the MM BM niche. Specialized CAR-like BMSCs in the niche secrete high levels of CXCL12, including CXCL12α and CXCL12γ. CXCL12α shows a relatively low affinity to HSPG and, upon secretion by BMSCs, will create a chemo-attractive gradient attracting MM cells to the niche. By contrast, CXCL12γ, having an extremely high affinity for HSPG, will be immobilized by HSPGs on the cell surface of BMSCs, inducing MM adhesion to the BMSCs. This CXCL12γ-controlled adhesion serves to retain MM cells in close physical contact with the BMSCs, providing MM cells with growth and survival signals through integrin receptors as well as with access to short-range growth and survival factors. This HSPG-immobilized CXCL12γ plays an important role in MM cell retention in the BM niche as well as in cell adhesion-mediated drug (CAM-DR) resistance in MM.

## Discussion

The CXCL12/CXCR4 axis plays a key role in the homing of normal plasma cell precursors and MM cells to the BM^9,10^, but the expression and specific role of CXCL12γ, a recently characterized CXCL12 isoform, which binds HSPGs with an extremely high affinity, has not been addressed. Here, we show that CXCL12γ is expressed *in situ* by reticular stromal cells in the human bone marrow niche as well as by BMSC lines and primary BMSC isolates. Unlike CXCL12α, CXCL12γ is immobilized on the cell surface of BMSCs by HSPGs, upon secretion. Functionally, this membrane bound CXCL12γ promotes adhesion of MM cells to the stromal niche cells, thereby protecting MM cells from drug-induced cell death.

Our study of the *in situ* expression of CXCL12γ shows that it is expressed by CAR-like reticular stromal cells in the BM. In normal BM, distinct CXCL12γ expression was present on stromal cells with long cytoplasmic processes, scattered among hematopoietic cells, as well as around adipocytes and capillaries, and in the endosteal zone (Figure 1A), areas with putative niche functions^32–37^. In BM sections of MM patients, CXCL12γ was also observed on stromal cells in areas infiltrated by MM cells. Notably, employing an antibody against an epitope shared by all CXCL12 isoforms, Abe-Suzuki *et al*^36^ recently reported a similar expression pattern, which also resembles the distribution of CAR cells in mouse bone marrow^9^. Study of isolated primary BMSCs and BMSC lines corroborates these findings, demonstrating that CXCL12γ is specifically expressed by isolated primary BMSCs and BMSC lines (Figure 1B).

CXCL12γ possesses an extraordinarily high affinity for HSPGs due to its unique C-terminal domain^15,20^. Interestingly, we observed that both primary BMCSs and HS5 cells constitutively express CXCL12γ on their cell surface, suggesting that this chemokine is retained by HSPGs upon secretion (Figure 1C). Indeed, we observed that KO of the HS-chain co-polymerase *EXT1* in HS5 BMSCs results in a complete loss of membrane-bound CXCL12γ. Importantly, immobilization by cell-surface HS was a specific feature of the CXCL12γ isoform, since overexpression of CXCL12α in HS5 did not result in detectable membrane retention, notwithstanding substantial intracellular expression (Figure 2).

We observed that specific deletion of CXCL12γ strongly reduces the capacity of HS5 BMSCs to mediate adhesion of MM cells to their cell surface. This result extends the previous observation that a total (*i.e.* non-isoform-specific) knockdown of CXCL12 reduces the capacity of BMSCs to mediate adhesion of MM cells^8^, pinpointing this effect to the CXCL12γ isoform. Similar to CXCL12γ deletion, *EXT1* deletion also attenuated MM cell adhesion to the BMSCs. Importantly, whereas the defective adhesion to HS5-CXCL12γKO cells could be overcome by exogenous expression of CXCL12γ, this could not correct the adhesion defect in HS5-EXT1KO cells, indicating that CXCL12γ immobilization by HSPGs is critically required (Figure 5). In line with this notion, in experiments employing recombinant CXCL12 to induce MM cell adhesion to VCAM-1 plastic, we observed that only immobilized (*i.e.* coated) CXCL12 effectively induced adhesion (Figure 3B).

Interaction of MM cells with BMSCs plays a central role in MM cell homing/retention and can also confer drug resistance^1,7^. We observed that co-culture with HS5 BMSCs of the HMCLs XG1 and MM1.S, and of primary MM cells did hardly or not affect tumor cell viability *per se,* but significantly reduced their sensitivity to the proteasome inhibitors bortezomib and carfilzomib. Interestingly, this resistance was largely annulled by specific deletion of CXCL12γ in BMSCs, identifying CXCL12γ as a major factor in the BMSC-mediated drug resistance. HS5 BMSCs cells with a deletion of *EXT-1* showed a similarly reduced capacity to protect MM cells, showing the essential role of membrane retention of CXCL12γ by HSPGs (Figure 6).

Drug resistance mediated by BMSCs can either be caused by soluble factors or by interactions via cell-adhesion molecules^1,42,43^. We observed that the protective effect of BMSCs to MM cells was largely abolished by physical separation of the MM and BMSCs, implying that this protection requires direct cell-cell contact (Figure 7A, B). This suggests that BMSCs might convey MM drug resistance via direct integrin-mediated signals, rather than by soluble growth and survival factors, although such factors are abundantly expressed by BMSCs^29,41,44^. However, recombinant CXCL12γ (or CXCL12α) induced adhesion to VCAM-1-coated plastic did not protect MM cells against bortezomib-induced cell death (Supplement Figure 6), indicating that integrin-mediated cell adhesion *per se* is not sufficient to instigate bortezomib resistance. Conceivably, CXCL12γ-controlled adhesion serves to retain MM cells in close physical contact with the BMSCs, providing MM cells with growth and survival signals through integrin receptors as well as with access to short-range growth and survival factors, such as Wnts and vascular endothelial growth factor^45,46^, which may act in concert to mediate drug resistance.

Our data suggest targeting CXCL12γ and/or its interaction with HSPGs, as a potential therapeutic strategy. Notably, MM cells express high levels of the HSPG syndecan-1, which is crucial for MM cell survival^47,48^ and promotes Wnt-mediated cell proliferation^29^ as well as hepatocyte growth factor (HGF), FGF, epidermal growth factor (EGF), and a proliferation-inducing ligand (APRIL) mediated signaling^49–51^. Hence, targeting HSPGs or the HS-biosynthesis machinery may disconnect the interaction of MM cells with the BM microenvironment at various levels^52^. Our studies corroborate previous studies, showing that disruption of the interaction between MM cells and BMSCs by the CXCR4 inhibitor AMD3100 enhances MM sensitivity to multiple therapeutic agents such as bortezomib, dexamethasone and melphalan^1,7,41^. Furthermore, targeting pan-CXCL12 by olaptesed pegol (ola-PEG), which neutralizes CXCL12 irrespective of the isoform, prevented MM progression in a murine model^8^, while a recent phase IIa clinical trial showed that patients with relapsed/refractory MM respond favorably to a combination of bortezomib or dexamethasone with ola-PEG^53^. Apart from CXCR4, MM cells also express CXCR7, an alternative receptor of CXCL12, which may also be involved in CAM-DR in MM^7^ as well as in MM progression^54^. Targeting the ligand CXCL12(γ) will simultaneously inhibit signaling through both chemokine receptors.

Taken together, our data suggest a scenario in which CXCL12γ functions as a ‘niche chemokine’ that, in conjunction with HSPGs, plays a key role in controlling adhesion, BM retention, and CAM-DR of MM cells (Figure 7C). These findings identify this unique membrane-bound chemokine, and associated HSPGs, as potential therapeutic targets in MM.

## Reference List

1. Azab AK, Runnels JM, Pitsillides C, et al. CXCR4 inhibitor AMD3100 disrupts the interaction of multiple myeloma cells with the bone marrow microenvironment and enhances their sensitivity to therapy. Blood. 2009;113(18):4341–4351.

2. Hideshima T, Mitsiades C, Tonon G, Richardson PG, Anderson KC. Understanding multiple myeloma pathogenesis in the bone marrow to identify new therapeutic targets. Nat Rev Cancer. 2007;7(8):585–598.

3. Pagnucco G, Cardinale G, Gervasi F. Targeting multiple myeloma cells and their bone marrow microenvironment. Ann N Y Acad Sci. 2004;1028:390–399.

4. Gay F, Magarotto V, Crippa C, et al. Bortezomib induction, reduced-intensity transplantation, and lenalidomide consolidation-maintenance for myeloma: updated results. Blood. 2013;122(8):1376–1383.

5. Engelhardt M, Terpos E, Kleber M, et al. European Myeloma Network recommendations on the evaluation and treatment of newly diagnosed patients with multiple myeloma. Haematologica. 2014;99(2):232–242.

6. van de Donk NW, Moreau P, Plesner T, et al. Clinical efficacy and management of monoclonal antibodies targeting CD38 and SLAMF7 in multiple myeloma. Blood. 2016;127(6):681–695.

7. Waldschmidt JM, Simon A, Wider D, et al. CXCL12 and CXCR7 are relevant targets to reverse cell adhesion-mediated drug resistance in multiple myeloma. Br J Haematol. 2017;179(1):36–49.

8. Roccaro AM, Sacco A, Purschke WG, et al. SDF-1 inhibition targets the bone marrow niche for cancer therapy. Cell Rep. 2014;9(1):118–128.

9. Sugiyama T, Kohara H, Noda M, Nagasawa T. Maintenance of the hematopoietic stem cell pool by CXCL12-CXCR4 chemokine signaling in bone marrow stromal cell niches. Immunity. 2006;25(6):977–988.

10. Peled A, Petit I, Kollet O, et al. Dependence of human stem cell engraftment and repopulation of NOD/SCID mice on CXCR4. Science. 1999;283(5403):845–848.

11. Sanz-Rodriguez F, Hidalgo A, Teixido J. Chemokine stromal cell-derived factor-1alpha modulates VLA-4 integrin-mediated multiple myeloma cell adhesion to CS-1/fibronectin and VCAM-1. Blood. 2001;97(2):346–351.

12. Alsayed Y, Ngo H, Runnels J, et al. Mechanisms of regulation of CXCR4/SDF-1 (CXCL12)-dependent migration and homing in multiple myeloma. Blood. 2007;109(7):2708–2717.

13. Menu E, Asosingh K, Indraccolo S, et al. The involvement of stromal derived factor 1alpha in homing and progression of multiple myeloma in the 5TMM model. Haematologica. 2006;91(5):605–612.

14. Yu L, Cecil J, Peng SB, et al. Identification and expression of novel isoforms of human stromal cell-derived factor 1. Gene. 2006;374:174–179.

15. Rueda P, Balabanian K, Lagane B, et al. The CXCL12gamma chemokine displays unprecedented structural and functional properties that make it a paradigm of chemoattractant proteins. PLoS One. 2008;3(7):e2543.

16. Zou YR, Kottmann AH, Kuroda M, Taniuchi I, Littman DR. Function of the chemokine receptor CXCR4 in haematopoiesis and in cerebellar development. Nature. 1998;393(6685):595–599.

17. Zhu W, Liang G, Huang Z, Doty SB, Boskey AL. Conditional inactivation of the CXCR4 receptor in osteoprecursors reduces postnatal bone formation due to impaired osteoblast development. J Biol Chem. 2011;286(30):26794–26805.

18. Nagasawa T, Hirota S, Tachibana K, et al. Defects of B-cell lymphopoiesis and bone-marrow myelopoiesis in mice lacking the CXC chemokine PBSF/SDF-1. Nature. 1996;382(6592):635–638.

19. Takabatake Y, Sugiyama T, Kohara H, et al. The CXCL12 (SDF-1)/CXCR4 axis is essential for the development of renal vasculature. J Am Soc Nephrol. 2009;20(8):1714–1723.

20. Laguri C, Sadir R, Rueda P, et al. The novel CXCL12gamma isoform encodes an unstructured cationic domain which regulates bioactivity and interaction with both glycosaminoglycans and CXCR4. PLoS One. 2007;2(10):e1110.

21. Rueda P, Richart A, Recalde A, et al. Homeostatic and tissue reparation defaults in mice carrying selective genetic invalidation of CXCL12/proteoglycan interactions. Circulation. 2012;126(15):1882–1895.

22. Esko JD, Selleck SB. Order out of chaos: assembly of ligand binding sites in heparan sulfate. Annu Rev Biochem. 2002;71:435–471.

23. Hacker U, Nybakken K, Perrimon N. Heparan sulphate proteoglycans: the sweet side of development. Nat Rev Mol Cell Biol. 2005;6(7):530–541.

24. Xu D, Esko JD. Demystifying heparan sulfate-protein interactions. Annu Rev Biochem. 2014;83:129–157.

25. Reijmers RM, Spaargaren M, Pals ST. Heparan sulfate proteoglycans in the control of B cell development and the pathogenesis of multiple myeloma. Febs j. 2013;280(10):2180–2193.

26. Ramani VC, Purushothaman A, Stewart MD, et al. The heparanase/syndecan-1 axis in cancer: mechanisms and therapies. Febs j. 2013;280(10):2294–2306.

27. Guo Z, Wang Z. The glypican Dally is required in the niche for the maintenance of germline stem cells and short-range BMP signaling in the Drosophila ovary. Development. 2009;136(21):3627–3635.

28. Pennetier D, Oyallon J, Morin-Poulard I, Dejean S, Vincent A, Crozatier M. Size control of the Drosophila hematopoietic niche by bone morphogenetic protein signaling reveals parallels with mammals. Proc Natl Acad Sci U S A. 2012;109(9):3389–3394.

29. Ren Z, van Andel H, de Lau W, et al. Syndecan-1 promotes Wnt/beta-catenin signaling in multiple myeloma by presenting Wnts and R-spondins. Blood. 2018;131(9):982–994.

30. Heckl D, Kowalczyk MS, Yudovich D, et al. Generation of mouse models of myeloid malignancy with combinatorial genetic lesions using CRISPR-Cas9 genome editing. Nat Biotechnol. 2014;32(9):941–946.

31. de Rooij MF, Kuil A, Geest CR, et al. The clinically active BTK inhibitor PCI-32765 targets B-cell receptor- and chemokine-controlled adhesion and migration in chronic lymphocytic leukemia. Blood. 2012;119(11):2590–2594.

32. Bonomo A, Monteiro AC, Goncalves-Silva T, Cordeiro-Spinetti E, Galvani RG, Balduino A. A T Cell View of the Bone Marrow. Front Immunol. 2016;7:184.

33. Morrison SJ, Scadden DT. The bone marrow niche for haematopoietic stem cells. Nature. 2014;505(7483):327–334.

34. Ribatti D, Basile A, Ruggieri S, Vacca A. Bone marrow vascular niche and the control of angiogenesis in multiple myeloma. Front Biosci (Landmark Ed). 2014;19:304–311.

35. Ribatti D, Nico B, Vacca A. Multiple myeloma as a model for the role of bone marrow niches in the control of angiogenesis. Int Rev Cell Mol Biol. 2015;314:259–282.

36. Abe-Suzuki S, Kurata M, Abe S, et al. CXCL12+ stromal cells as bone marrow niche for CD34+ hematopoietic cells and their association with disease progression in myelodysplastic syndromes. Lab Invest. 2014;94(11):1212–1223.

37. Omatsu Y, Sugiyama T, Kohara H, et al. The essential functions of adipo-osteogenic progenitors as the hematopoietic stem and progenitor cell niche. Immunity. 2010;33(3):387–399.

38. Azab AK, Azab F, Blotta S, et al. RhoA and Rac1 GTPases play major and differential roles in stromal cell-derived factor-1-induced cell adhesion and chemotaxis in multiple myeloma. Blood. 2009;114(3):619–629.

39. Connell BJ, Sadir R, Baleux F, et al. Heparan sulfate differentially controls CXCL12alpha- and CXCL12gamma-mediated cell migration through differential presentation to their receptor CXCR4. Sci Signal. 2016;9(452):ra107.

40. Di Marzo L, Desantis V, Solimando AG, et al. Microenvironment drug resistance in multiple myeloma: emerging new players. Oncotarget. 2016;7(37):60698–60711.

41. Ghobrial IM, Detappe A, Anderson KC, Steensma DP. The bone-marrow niche in MDS and MGUS: implications for AML and MM. Nat Rev Clin Oncol. 2018;15(4):219–233.

42. Gupta D, Treon SP, Shima Y, et al. Adherence of multiple myeloma cells to bone marrow stromal cells upregulates vascular endothelial growth factor secretion: therapeutic applications. Leukemia. 2001;15(12):1950–1961.

43. Hideshima T, Nakamura N, Chauhan D, Anderson KC. Biologic sequelae of interleukin-6 induced PI3-K/Akt signaling in multiple myeloma. Oncogene. 2001;20(42):5991–6000.

44. Rougier F, Cornu E, Praloran V, Denizot Y. IL-6 and IL-8 production by human bone marrow stromal cells. Cytokine. 1998;10(2):93–97.

45. Farin HF, Jordens I, Mosa MH, et al. Visualization of a short-range Wnt gradient in the intestinal stem-cell niche. Nature. 2016;530(7590):340–343.

46. Barkefors I, Le Jan S, Jakobsson L, et al. Endothelial cell migration in stable gradients of vascular endothelial growth factor A and fibroblast growth factor 2: effects on chemotaxis and chemokinesis. J Biol Chem. 2008;283(20):13905–13912.

47. Yang Y, MacLeod V, Dai Y, et al. The syndecan-1 heparan sulfate proteoglycan is a viable target for myeloma therapy. Blood. 2007;110(6):2041–2048.

48. Reijmers RM, Groen RW, Rozemuller H, et al. Targeting EXT1 reveals a crucial role for heparan sulfate in the growth of multiple myeloma. Blood. 2010;115(3):601–604.

49. Derksen PW, Keehnen RM, Evers LM, van Oers MH, Spaargaren M, Pals ST. Cell surface proteoglycan syndecan-1 mediates hepatocyte growth factor binding and promotes Met signaling in multiple myeloma. Blood. 2002;99(4):1405–1410.

50. Mahtouk K, Cremer FW, Reme T, et al. Heparan sulphate proteoglycans are essential for the myeloma cell growth activity of EGF-family ligands in multiple myeloma. Oncogene. 2006;25(54):7180–7191.

51. Reijmers RM, Groen RW, Kuil A, et al. Disruption of heparan sulfate proteoglycan conformation perturbs B-cell maturation and APRIL-mediated plasma cell survival. Blood. 2011;117(23):6162–6171.

52. Sanderson RD, Yang Y. Syndecan-1: a dynamic regulator of the myeloma microenvironment. Clin Exp Metastasis. 2008;25(2):149–159.

53. Ludwig H, Weisel K, Petrucci MT, et al. Olaptesed pegol, an anti-CXCL12/SDF-1 Spiegelmer, alone and with bortezomib-dexamethasone in relapsed/refractory multiple myeloma: a Phase IIa Study. Leukemia. 2017;31(4):997–1000.

54. Azab AK, Sahin I, Moschetta M, et al. CXCR7-dependent angiogenic mononuclear cell trafficking regulates tumor progression in multiple myeloma. Blood. 2014;124(12):1905–1914.

