## Supplemental material and data for "The CXCL12gamma chemokine immobilized by heparan sulfate on stromal niche cells controls adhesion and mediates drug resistance in multiple myeloma"

#### Supplemental table 1: qPCR primer sequence

| | |
|---|---|
| CXCL12α forward | CCAAACTGTGCCCTTCAGAT |
| CXCL12α reverse | CGTCTTTGCCCTTTCATCTC |
| CXCL12γ forward | CCAAACTGTGCCCTTCAGAT |
| CXCL12γ reverse | CTTTTCTGGGCAGCCTTTCT |
| RPLPO forward | GCTTCCTGGAGGGTGTCCGC |
| RPLPO reverse | TCCGTCTCCACAGACAAGGCCA |

**Supplemental figure legends**

**Figure 1.** CXCL12α mRNA expression in HS5-WT and HS5-CXCL12γ KO cells. A representative for 2 independent experiments. ns, $P > 0.05$ using unpaired student's t-test.

**Figure 2.** MTT assay analysis of the growth of HS5-WT, HS5-CXCL12γKO and HS5-EXT1KO cells. Mean ±SD of three independent experiments in triplicate is shown.

**Figure 3. (A)** Adhesion of the HMCL L363 to HS5-WT and two HS5-CXCL12γKO clones. Mean ±SD of three independent experiments in triplicate. *, $P \leq 0.05$ using one-way ANOVA analysis. **(B)** Adhesion of the HMCL L363 to HS5-WT and HS5-EXT1KO. Mean ±SD of 3 independent experiments in triplicate. *, $P \leq 0.05$ using unpaired student's t-test.

**Figure 4. (A)** Flow cytometric discrimination of MM cells from BMSCs expressing GFP. **(B)** BMSC viability is not affected by bortezomib. HS5 cells were co-cultured with XG1 cells in the presence of bortezomib at indicated concentrations for 3 days. HS5 viability was analyzed by flow cytometry.

**Figure 5.** CXCL12γ mediates carfilzomib resistance in XG1 cells. XG1 was cultured alone, or co-cultured with HS5-WT, HS5-CXCL12γKO or HS5-EXT1KO, in the presence carfilzomib for 3 days. Mean ±SD of 3 independent experiments in triplicate. *, $P \leq 0.05$; **, $P \leq 0.01$ using one-way ANOVA analysis.

**Figure 6.** Recombinant CXCL12γ (or CXCL12α) induced adhesion to VCAM-1 does not protect HMCLs from bortezomib-induced cell death. The HMCLs XG1 or MM1.S were cultured on a surface co-coated with CXCL12γ or CXCL12α and VCAM-1 in the presence of bortezomib for 3 days. Cell viability was analyzed by flow cytometry. A representative plot for 2 independent experiment is shown.

### Supplemental data

#### Figure 1

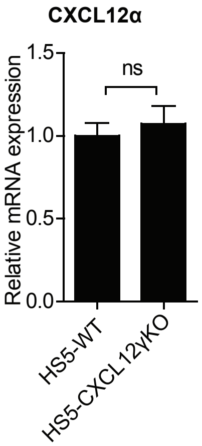


#### Figure 2

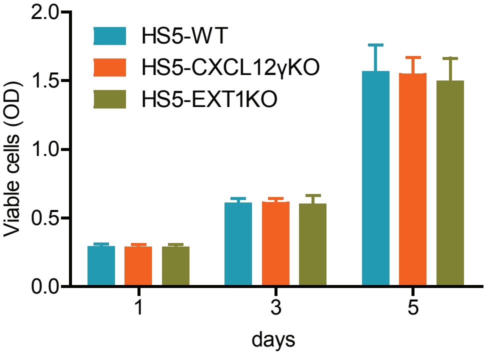


#### Figure 3

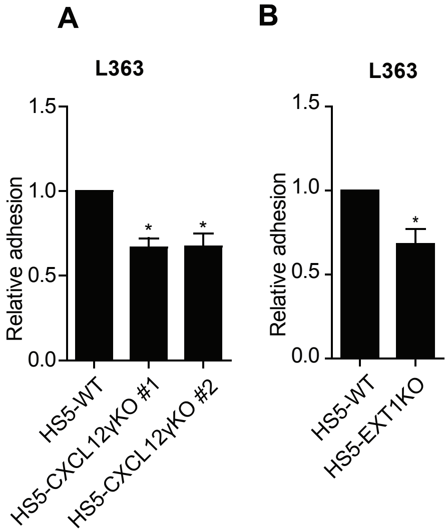


#### Figure 4

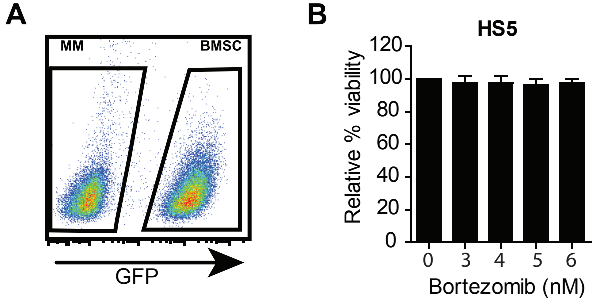


#### Figure 5

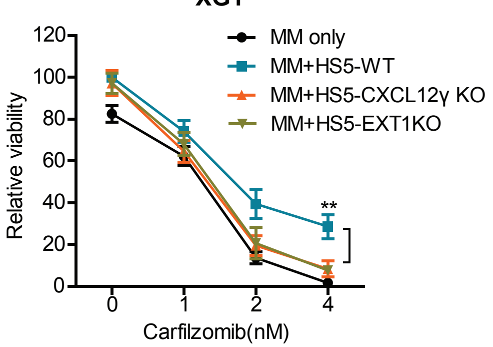


#### Figure 6

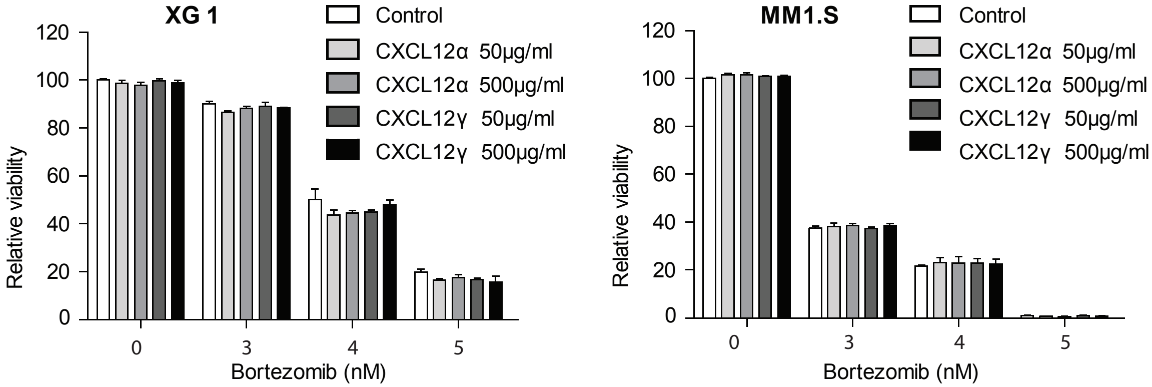
